# Word-selective EEG/MEG responses in the English language obtained with Fast Periodic Visual Stimulation (FPVS)

**DOI:** 10.1101/2024.08.02.606349

**Authors:** O Hauk, M Marchive, A Volfart, C Schiltz, B Rossion, MA Lambon Ralph, A Lochy

## Abstract

Fast periodic visual stimulation (FPVS) allows the recording of objective brain responses of human word discrimination (i.e., reproducible word-category-selective responses) with a high signal-to-noise ratio. This approach has been successfully employed over the last decade in a number of scalp electroencephalography (EEG) studies. Three important advances were achieved in this study: (1) robust measures of written word-selective responses with this approach have not been reported in English; (2) responses have only been reported in EEG but not with MEG, and (3) without source localization. Thus, we presented English words periodically (2 Hz) among different types of letter strings (10Hz; consonant strings, non-words, pseudowords) whilst recording simultaneous EEG and MEG in 25 participants who performed a simple non-linguistic color detection task. Data were analyzed in sensor and in source space (L2-minimum-norm estimation, MNE). With only 4 minutes of stimulation we observed a robust word discrimination response in each condition including, importantly, even when words were embedded in word-like pseudowords. This response was larger in nonwords and largest in consonant strings. We observed left-lateralized responses in all conditions in the majority of our participants. Cluster-based permutation tests revealed that these responses were left-lateralized in sensor as well as in source space, with peaks in left posterior regions. Our results demonstrate that the FPVS paradigm can elicit robust English word-discrimination responses in EEG and MEG within only a few minutes of recording time. Together with source estimation, this can provide novel insights into the neural basis of visual word recognition in healthy and clinical populations.

## Introduction

Recognizing written words at a single glance is a crucial marker of reading expertise. Most adults in literate societies read fluently and effortlessly, often unaware of how complex this ability is at the cognitive level. In natural reading, a normal reader can discriminate and identify words at a rate of several words per second, offering the opportunity to present a large number of stimuli in a short amount of time. Here, we used a fast periodic visual stimulation (FPVS) paradigm to study EEG/MEG brain responses that discriminate between words and different types of non-word letter strings.

Over the past decade, FPVS-EEG has been used to investigate visual word recognition processes in adults (e.g. Lochy et al., 2018; Lochy, Van Belle, & Rossion, 2014, 2015) or in developmental studies (de Ghelcke, Rossion, Schiltz, & Lochy, 2020, 2021; Lutz, Coraj, Fraga-González, & Brem, 2024; Wang et al., 2024; Wang et al., 2023). Coupled with electroencephalography (EEG), FPVS measures word-selective neural responses over or in the left occipito-temporal cortex implicitly, objectively and with a high signal-to-noise ratio (SNR) (Lochy et al., 2018; Lochy et al., 2015). In the so-called oddball version of the paradigm, words are presented periodically (e.g., every five items) as deviants (oddballs) in streams of base stimuli (e.g., pseudowords) displayed at a certain frequency (e.g., 10Hz). If words are discriminated by the visual word recognition system from the base stimuli, then a word-selective response occurs exactly at the predefined oddball frequency (e.g., in base stimuli displayed at 10Hz, words occur at 10/5Hz, and responses are detectable at 2Hz (Lochy et al., 2024; Lochy et al., 2015)). Importantly, the word-selective response represents a measure of differential processing between base and oddball stimuli, and therefore in the case of words among pseudowords it reflects the neural response to words over and above the responses that words and pseudowords have in common. Thus similar to the logic used in some behavioural studies of lexical decision (e.g. Evans, Lambon Ralph, & Woollams, 2012), the oddball response is constrained by the contrast experimentally defined between the two stimulus categories, and provides a measure of coarse discrimination when words are inserted among pseudo-letters, while fine-grained prelexical or lexical-semantic discrimination is measured when words appear among nonwords or pseudowords (Lochy et al., 2015).

In spite of these clear observations and the significant advantages of the approach, its extension to English remains uncommon and sometimes elusive. In addition to the dearth of FPVS word recognition studies in general, one recent study with ten participants encountered difficulties replicating the significant results of the original French paradigm in English (Barnes, Petit, Badcock, Whyte, & Woolgar, 2021). This could be due to linguistic differences between French and English, or to different methodological choices (see Lochy et al., (2024), and discussion of the present study). Therefore, the application of FPVS to English word recognition needs further development before it can serve as an individually-sensitive measure of written word processing in basic and clinical research. This was the goal of this major study.

Our study also addressed two other important targets: to date word-selective responses have only been reported in EEG but not MEG, and without source estimation. Possible future applications of the word-selective FPVS paradigm include the fast determination of lateralized brain responses to linguistic stimuli. However, it is well-known that sensor-space signals are difficult, if not impossible, to relate to putative brain regions without very strong prior information, in particular for EEG (e.g. Ahlfors et al., 2010). The highest spatial resolution can be obtained using a combination of EEG and MEG recordings, as they are sensitive to different spatial aspects of the neural current distributions (Ahlfors et al., 2010; Goldenholz et al., 2009; Hauk, Stenroos, & Treder, 2019; Henson, Mouchlianitis, & Friston, 2009; Molins, Stufflebeam, Brown, & Hamalainen, 2008). Thus, we evaluated the relative sensitivities of EEG and MEG sensor types as well as source estimates to word-selective FPVS responses using English stimulus material, particularly with respect to hemispheric laterality.

## Methods

### Participants

Twenty-nine participants were initially recruited from the MRC CBU’s volunteer panel database. Data from four of them had to be excluded because of technical problems or high noise levels, leaving 25 datasets for the final analysis. Of those, 14 identified as females. Mean age was 27 years (SD 6 years, range 19-40 years). All participants reported to be right-handed native English speakers, to have normal or corrected-to-normal vision, and to have no history of neurological or developmental disorders. They were monetarily reimbursed for their participation to the study. This study was approved by the Cambridge Psychology Research Ethics Committee.

### Stimuli

Our stimuli and experimental settings were close to those described in Lochy et al. (2024). The stimuli consisted of 30 English high frequency nouns composed of five letters (frequency mean 81.79 per million ± 139.57) with an average of 3.4 (±2.69) orthographic neighbours and an average bigram frequency of 10335 (±3.259) calculated with Wordgen (Duyck, Desmet, Verbeke, & Brysbaert, 2004). 120 consonant strings were created by randomly mixing consonant strings. Additionally, we took each of the 120 words and constructed a five-letter pseudo-word with the same consonant/vowel structure, bigram frequency (9279 ± 2563) and number of orthographic neighbours (2.48 ± 1.85). These pseudo-words were pronounceable and respected the phonological rules of the English language, but did not correspond to real words. They did not differ to the words with respect to bigram frequency (U = 1424, p = 0.078 with Mann-Whitney correction) and orthographic neighbours (U = 1454, p = 0.100 with Mann-Whitney correction). A set of 120 non-words were created by shuffling the letters of the pseudo-words, hence containing five letters that were not pronounceable and did not follow the orthographic rules in English. Compared to words, and as expected, these non-words were significantly different in terms of bigram frequency (mean 4159 ± 2412) (t(148) = -11.637, p<.001) and orthographic neighbours (mean 0 ± 0).

All stimuli were presented on a grey background in a black Verdana font and varied randomly in size spanning 80% to 120% of their average size. They were shown on a screen at a distance of approximately 1.3 m in front of the participant, the stimuli subtended a maximum 4 degrees of visual angle.

**Figure 1:**
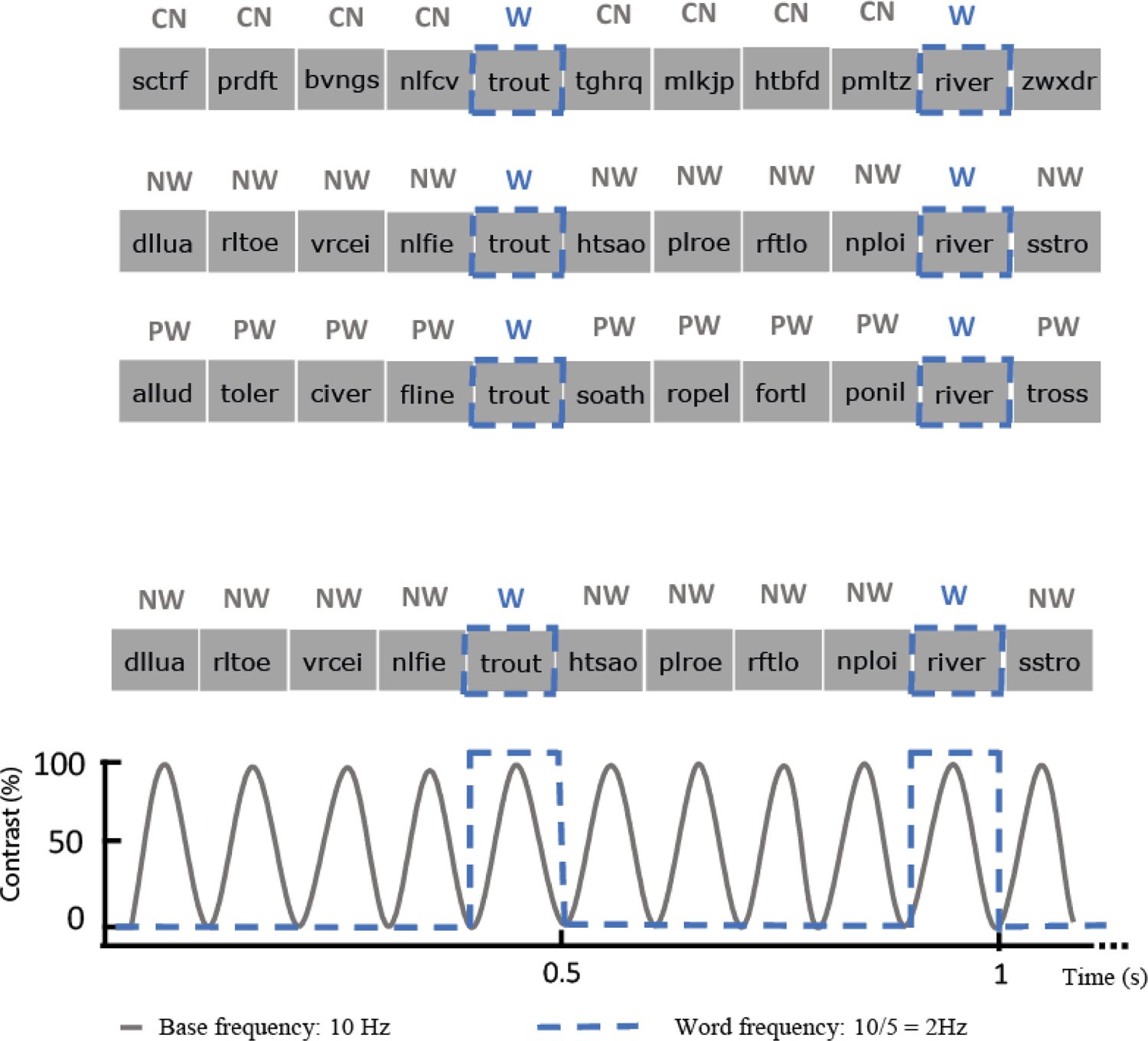
Illustration of the FPVS words paradigm. Consonant strings (CN), non-words (NW) or pseudo-words (PW) stimuli were presented at base frequency of 10 Hz using a sinusoidal contrast modulation. Words were inserted every 5 stimuli, corresponding to a word-selective frequency of 2Hz (= 10 Hz/5).

### Procedure

A schematic illustration of the FPVS paradigm is presented in **Figure 1**. Stimuli were presented to participants using Java 8. All stimuli were presented on a uniform grey background using sinusoidal contrast manipulation, from 0 to 100% to 0% for each stimulus (Rossion, Torfs, Jacques, & Liu-Shuang, 2015). Monitor refresh rate was 60 Hz. Participants completed four runs of letter-string stimulation for each contrast (i.e., a contrast with either consonant strings, nonwords or pseudowords) leading to a total of 12 runs. Each run consisted of letter strings presented at a base frequency of 10 Hz (100 ms per stimulus), and every fifth image was a word (“oddball” frequency 2 Hz). A neural response that does not differ systematically between words and other types of letter strings will project on the 10 Hz component of the EEG/MEG spectrum and its harmonics (20 Hz, 30 Hz, etc.), whereas a differential response for words and non-word stimuli (i.e., a word-selective response) will be reflected at 2 Hz and harmonics (4 Hz, 6 Hz, etc.). Each run lasted for 64 seconds, including 2 seconds of fade-in and fade-out at the beginning and at the end of the sequence, respectively.

As in previous studies (e.g. Rossion et al., 2015), participants were instructed to perform a colour change detection task during the stimulus presentation period. The stimuli were presented together with two lateral vertical bars that changed colour simultaneously or in isolation. Participants were asked to press a button with their right index finger when both bars changed from blue to red at the same time. This occurred, randomly, eight times per run and lasted for 200 ms. The minimum difference in time between each colour change was 2 seconds. This task was chosen because it is orthogonal to the experimental manipulation (i.e., different types of letter strings), and has recently been shown to produce larger word-selective responses than a previously used central cross colour-change detection task (Lochy et al., 2024). Its purpose is to ensure that participants pay attention to the stimuli. This was confirmed, as performance was high with an average correct target detection rate of 98% (standard deviation 3%, min|max 89%|100%) and an average response time of 459 ms (SD 66 ms, min|max 347ms|597ms).

### Data acquisition

EEG/MEG data were acquired on an Elekta Neuromag Triux neo system (Elekta AB, Stockholm, Sweden), containing 306 MEG sensors (102 magnetometers and 204 gradiometers), and 64 EEG electrodes mounted on an Easycap cap (EasyCap GmbH, Herrsching, Germany). The EEG recording reference electrode was attached to the nose, and the ground electrode to the left cheek. The electrooculogram (EOG) was recorded from electrodes above and below the left eye (vertical EOG) and at the outer canthi (horizontal EOG). The sampling rate during data acquisition was 1000 Hz and an on-line band pass filter 0.03 to 330 Hz was applied. Prior to the EEG/MEG recording, the positions of 5 Head Position Indicator (HPI) coils were attached to the EEG cap (for head localisation inside the scanner and continuous movement tracking), 3 anatomical landmark points (two preauricular points and nasion) as well as the EEG electrodes and about 50-100 additional points that cover most of the scalp were digitised using a 3Space Isotrak II System (Polhemus, Colchester, Vermont, USA) for later co-registration with MRI data. Our data can be made available via the MRC Cognition and Brain Sciences Unit’s data repository on request.

High-resolution structural T1-weighted MRI images were acquired in a 3T Siemens Tim Trio scanner at the MRC Cognition and Brain Sciences Unit (UK) with a 3-D magnetization prepared rapid gradient-echo sequence, field of view = 256 mm × 240mm × 192mm, matrix dimensions = 256 × 240 × 160, 1-mm isotropic resolution, repetition time = 2250 ms, inversion time = 900 ms, echo time = 2.99 ms, and flip angle = 9°.

### Sensor Space Analysis

#### Pre-processing

MEG data were subjected to spatio-temporal signal-space separation (SSS) implemented in the Maxfilter software (Version 2.2.12) of Elekta Neuromag to remove noise generated from sources distant to the sensor array (Taulu & Kajola, 2005; Taulu & Simola, 2006). The SSS procedure included movement compensation (locations recorded every 200 ms), bad MEG channel interpolation, and temporal SSS extension (with default buffer length 10 s and sub-space correlation limit 0.98). The origin in the head frame is chosen as (0,0,45) mm.

The following steps of analysis were performed in MNE-Python software package (Version 1.4.0, http://martinos.org/mne/stable/index.html) (Gramfort et al., 2013). After visual inspection of the raw data, bad EEG channels (determined by visual inspection) were interpolated. A notch filter at 50 and 100 Hz was then applied, followed by a low-pass filter with cut-off frequency 140 Hz. In order to remove eye movements and heart artefact an Independent Component Analysis (ICA) was computed, removing a maximum of 4 ICA components (2 for heart, 2 for eyes). The exact ICA procedure closely followed the examples provided for the MNE-Python software (https://martinos.org/mne/dev/auto_tutorials/plot_ica_from_raw.html), which uses the temporal correlation between ICA components and EOG channels as a criterion for the removal of ICA components.

#### Frequency- and time-domain analyses

First, data of the four 60-s runs (with fade-in and –out periods removed) were averaged in the time domain to improve SNR, and a Fast Fourier Transform (FFT) with frequency resolution 0.016 was applied. We divided the FFT spectrum into ten segments of +/-0.35 Hz centred at the frequency of interest and its nine higher harmonics, which included 21 frequency samples at each side of the centre frequency. These segments were summed up for ten harmonics. For the word-selective frequency, i.e. 2 Hz, multiples of the base frequency (i.e., 10 Hz and 20 Hz) were excluded. In order to correct for the variations in baseline noise levels around each frequency of interest, the amplitude of ten neighbouring frequency-bins on each side of the centre frequency was averaged and subtracted from each frequency bin (Retter, Jiang, Webster, & Rossion, 2020). A gap of one frequency bin on each side of the target frequency was included to avoid spectral leakage. The minimum and maximum values were also removed from the baseline interval. We applied the same summing and baseline correction procedure to the harmonics of the base frequency. Finally, Z-scores for word-selective and base frequencies were computed by dividing the baseline-corrected amplitudes by the standard deviation of the neighbouring bins from the baseline correction procedure.

Sensor space results are presented separately for the three sensor types employed in this study: EEG, gradiometers and magnetometers. The EEG measures the electric potential in Volts, magnetometers the magnetic flux in Tesla, and gradiometers the magnetic flux gradient in two orthogonal planar directions in Tesla per centimetre. As there are two gradient measurements per sensor location, the values of each gradiometer pair were plotted as the root-mean-square (RMS) per pair. Note that we present all results as Z-scores, which are comparable across sensor types.

We tested the reliability of our word-selective responses at the group level using cluster-based permutation testing, which is a common procedure to address the multiple-comparisons problem in EEG/MEG data (Maris & Oostenveld, 2007). The summed Z-score distributions across sensors at the centre frequency bin were subjected to two-tailed t-tests to first define clusters at a single-sensor p-value threshold of 0.05, and then the significance of clusters was established in a permutation procedure (10000 permutations) and with a cluster-p-threshold of 0.05.

The laterality of sensor-space responses was investigated by focussing on EEG for comparison with previous FPVS studies, as well as on gradiometers which allow a closer association between peaks in the distribution and the putative location of sources underneath those sensors than EEG and magnetometers (Hämäläinen, Hari, Ilmoniemi, Knuutila, & Lounasmaa, 1993). For EEG, we selected electrodes PO7 and PO8 in left and right posterior scalp regions, respectively, as there were the only electrodes that overlapped between our EEG setup in the MEG and the previous study by Lochy et al. (2015). Note the neighbouring EEG electrodes are sensitive to similar brain structures due to volume conduction. Since no comparable data from MEG studies were available, we chose pre-defined groups of left and right temporal gradiometers and used their respective averages (options “Left-temporal” and “Right-temporal” from MNE-Python’s function ‘read_vectorview_selection’).

### Source Space Analysis

Source estimation was performed on combined EEG/MEG data using L2-minimum-norm estimation, as appropriate for data where the number and location of sources is not known a-priori (Hämäläinen & Ilmoniemi, 1984; Hauk, 2004). In the frequency domain, source estimates were computed for the summed topographies across harmonics for the word-selective and base frequencies, respectively. This was performed in MNE-Python software with standard parameters settings. We used individual MRI images for head-modelling. The MRI data were pre-processed in Freesurfer V6.0.0 (Fischl, 2012), and the head model (3-layer boundary element model) created in MNE-Python. We used L2-minimum-norm estimation without depth-weighting or noise-normalisation, and with a loose orientation constraint (ratio of variances between tangential and normal dipole components: 0.2). We used a regularisation parameter based on a SNR value of 3 (default in the software). The procedures for baseline-correction and z-scoring in the frequency domains as described above were also applied to source-space results.

The significance of word-selective responses in source space was determined analogously to our procedure in sensor space using cluster-based permutation testing. Laterality in source space was examined by collapsing Z-scores across most of the temporal lobes, by combing the regions-of-interest (ROIs) inferior, middle and superior temporal lobe from the Desikan-Killiany cortical parcellation in the left and right hemisphere, respectively (https://surfer.nmr.mgh.harvard.edu/fswiki/CorticalParcellation).

## Results

Figure 2 shows the grand-averaged Z-scored frequency spectra for the condition in which words were interspersed among consonant strings. The peak at 10 Hz reflects the base frequency, i.e., the common response to all stimuli, while the peaks at the harmonics of 2 Hz represent word-selective response, which discriminates between words and consonant strings in this example. The topographies of Z-scores are shown for the first four harmonics of the word-selective response and for each sensor type at the top of each panel. All sensor types show reliable word-selective responses in posterior sensors.

**Figure 2:**
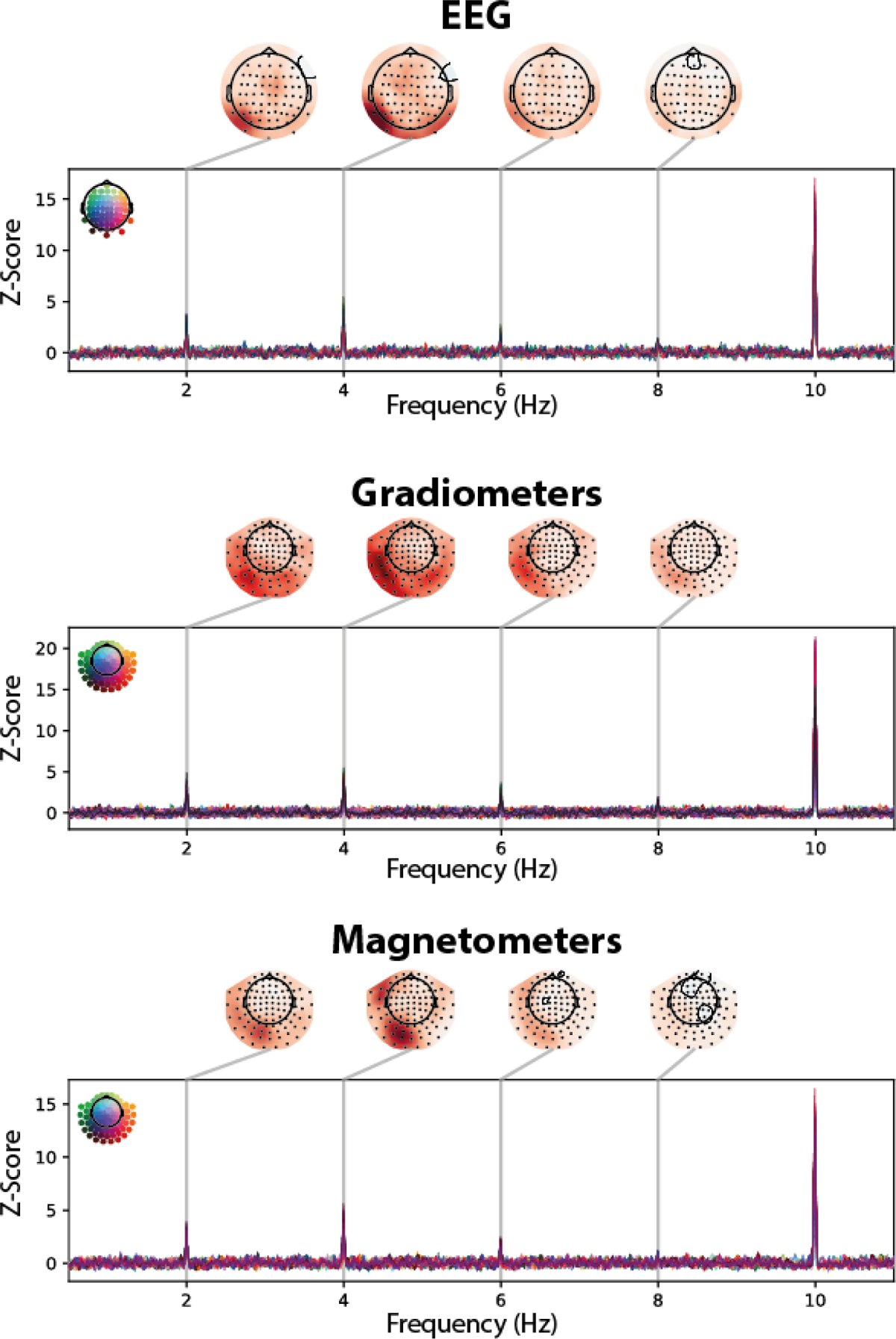
Z-scored spectra for the CN condition. Each panel shows the z-scored frequency spectra for individual channels types (EEG, gradiometers, magnetometers) used in our EEG/MEG recordings between 0.5 and 11Hz. The peak at 10Hz corresponds to the base frequency of visual stimulus presentation, and the peaks at multiples of 2 Hz correspond to responses at the word-selective frequency and its harmonics. Line colour indicates the position of the corresponding sensor in the sensor array (inlet in top left of panels). The topographies at the top of each panel represent Z-score distributions for the first four harmonics of the word-selective responses.

Figure 3 presents the Z-scored summed frequency spectra around harmonics of the word-selective frequencies for each condition (words among consonant strings, CN; words among non-words, NW; words among pseudowords, PW). For each condition, all three sensor types are shown (EEG, gradiometers, magnetometers). While Z-scores are highest in the CN and lowest in the PW condition, a peak at the centre frequency (0 Hz) above the baseline indicates a reliable word-selective response across participants in all conditions and sensor types. This is confirmed by the topographies on the right, which show the Z-score distributions for the summed spectra at the centre frequency. All topographies are characterised by left-lateralised peaks in posterior areas.

**Figure 3:**
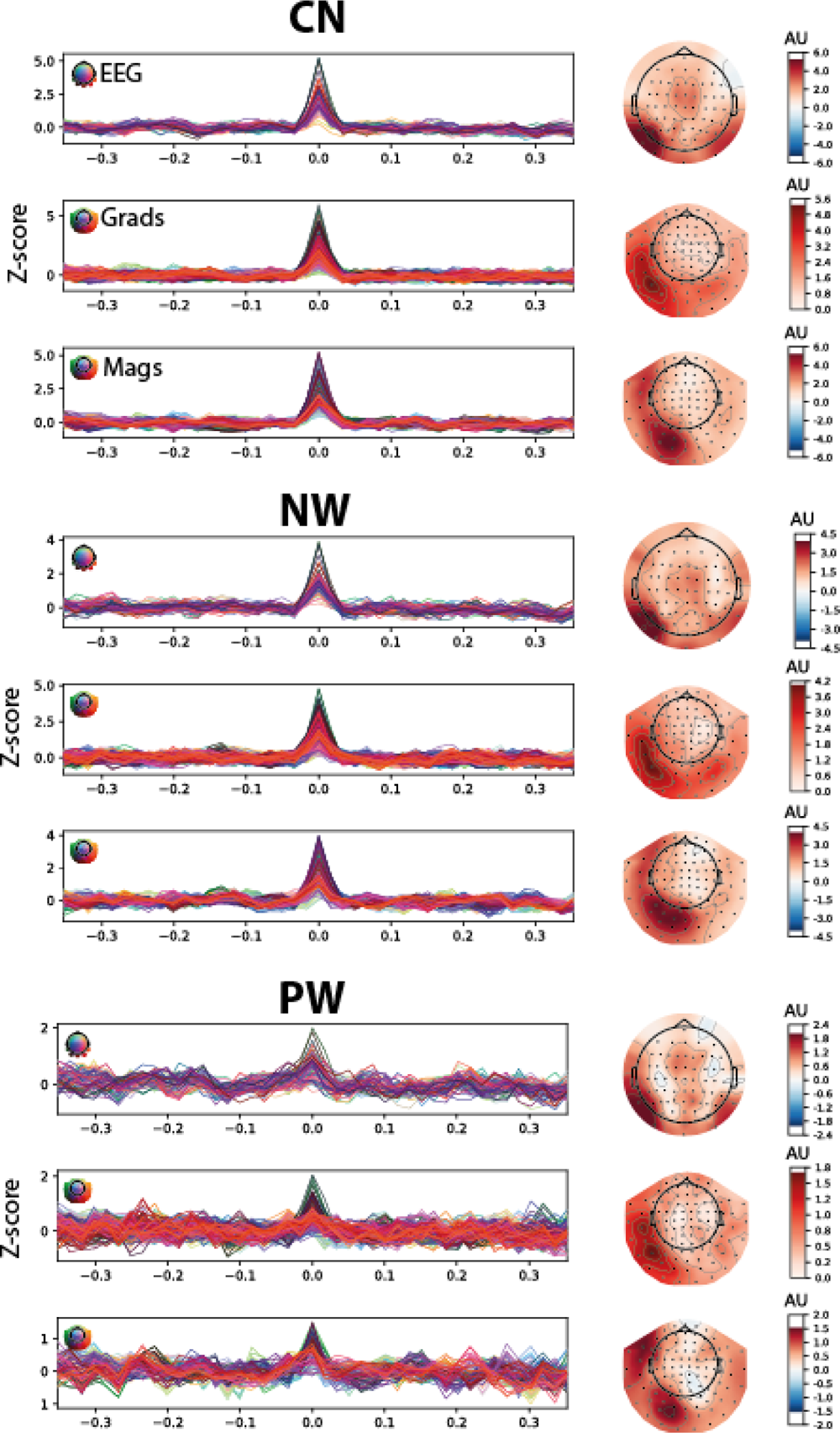
Z-scored summed frequency spectra around harmonics of word-selective frequencies. Each group of three panels presents results for one condition for all three sensor types. Each panel contains an overlay of all sensors (coloured lines) within EEG (top of each condition), gradiometers (“Grads”, middle) and magnetometers (“Mags”, bottom). The peak at 0 Hz for word-selective responses indicates reliable discrimination between words and pseudowords. Topographies on the right represent averaged z-score distributions at the centre frequency. CN: consonant strings; NW: nonwords; PW: pseudowords.

**Figure 4:**
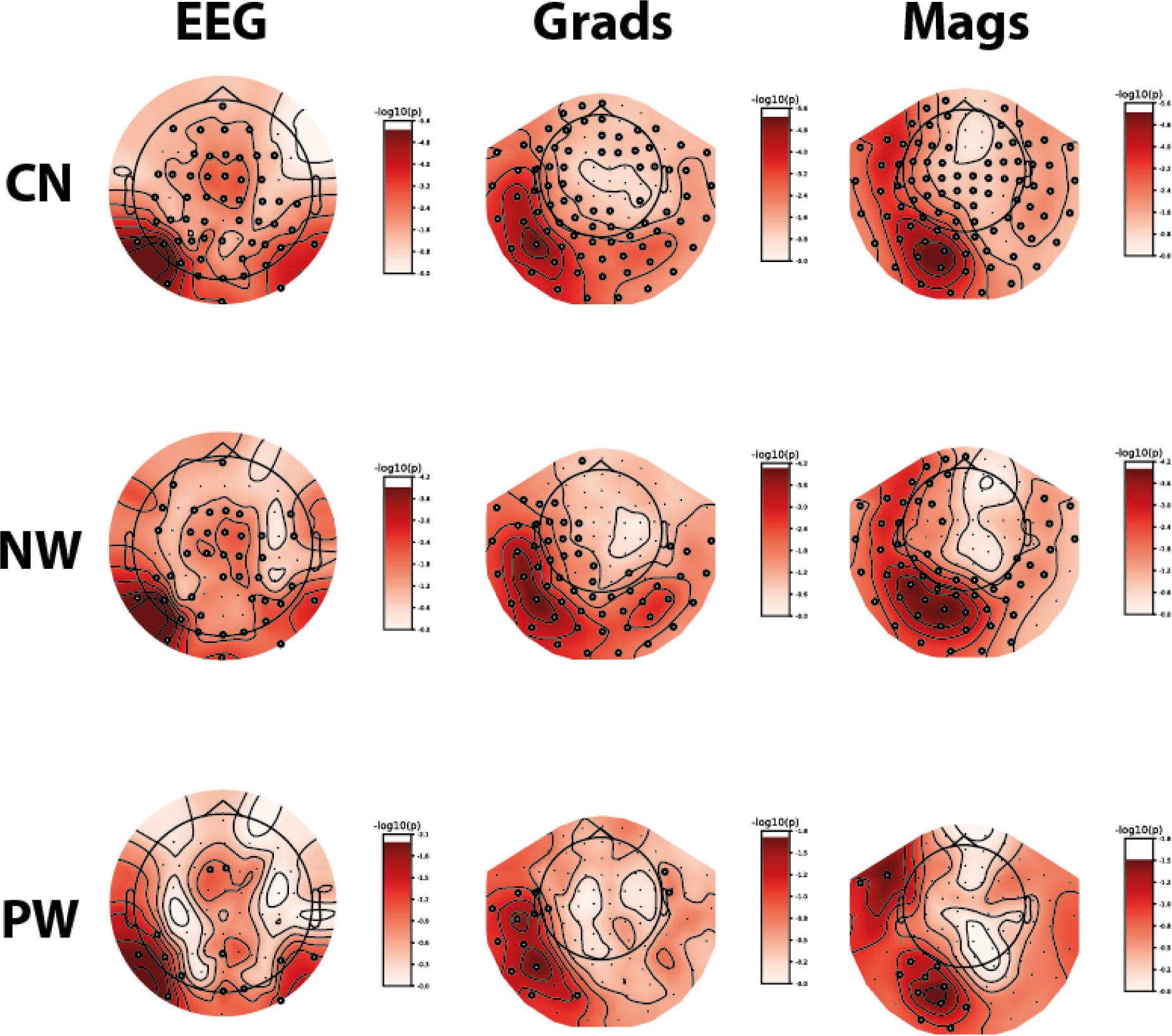
Statistics in sensor space. Summed Z-scored spectra were tested against zero using cluster-based permutation tests for each sensor type separately. Black circles indicate sensors that are part of a significant cluster. CN: consonant strings; NW: nonwords; PW: pseudowords.

Figure 4 displays the corresponding cluster-based permutation results in sensor space for each condition and sensor type. Small black circles indicate sensors that are part of significant clusters. While the number of significant channels decreases from CN over NW to PW, the dominant left-hemispheric peaks are significant in all cases.

This pattern of results is confirmed by our source space analysis in Figure 5. Here, Z-score distributions across the cortical surface obtained from L2-minimum-norm estimates were masked with the significant clusters from a cluster-based permutation test. Again, the extent of significant clusters follows the pattern CN>NW>PW, but several peaks in the left temporal lobe are significant in all conditions.

**Figure 5:**
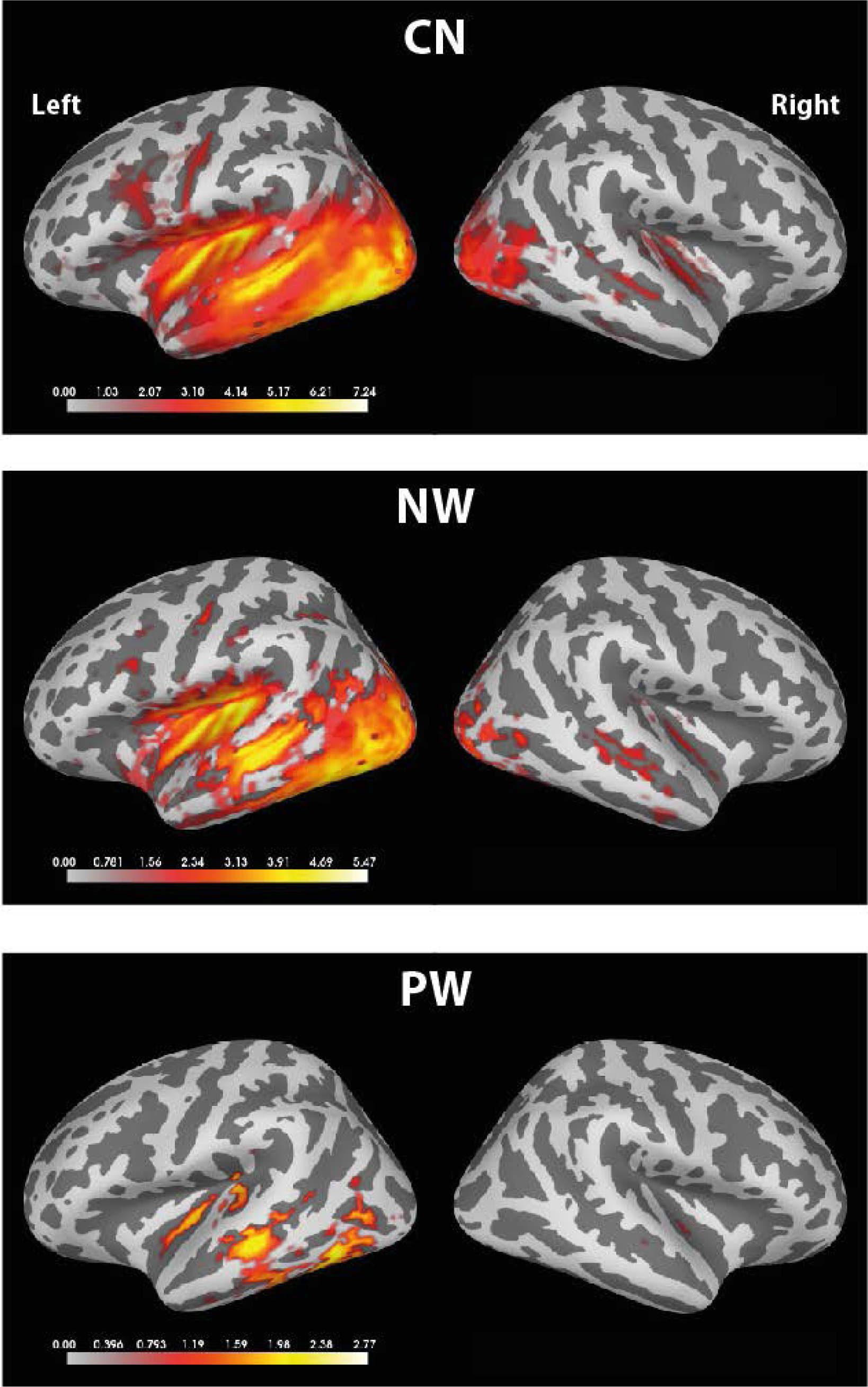
Source space analysis of word-specific responses. The panels show word-specific Z-scored summed spectra in source space (based on L2 minimum-norm estimation) masked by significant clusters from a cluster-based permutation test against zero. CN: consonant strings; NW: nonwords; PW: pseudowords.

**Table 1:** Statistical results for laterality of Z-scores. T-and p-values are shown for word-selective and base responses, respectively.

|  | **EEG**<br>word-selective \| base<br>t (p) | **Gradiometers**<br>word-selective \| base<br>t (p) | **Temporal Lobe**<br>word-selective \| base<br>t (p) |
| --- | --- | --- | --- |
| **CN** | 1.7 (0.11) \| -1.6 (0.13) | 7.0 (3.4e-7) \| -1.6 (0.13) | 6.4 (0.000001) \| -1.9 (0.06) |
| **NW** | 1.4 (0.17) \| -1.0 (0.33) | 5.6 (0.00001) \| -1.8 (0.08) | 4.2 (0.0003) \| 0.0 (0.99) |
| **PW** | 0.5 (0.62) \| -0.4 (0.71) | 3.4 (0.002) \| -1.5 (0.16) | 5.6 (0.000009) \| -1.0 (0.32) |

**Table 2:** Percentage of participants with left-lateralised FPVS responses in sensor and source space for word-selective and base responses, respectively.

|  | **EEG**<br>word-selective \| base | **Gradiometers**<br>word-selective \| base | **Temporal Lobe**<br>word-selective \| base |
| --- | --- | --- | --- |
| **CN** | 60% \| 44% | 96% \| 36% | 88% \| 44% |
| **NW** | 52% \| 52% | 88% \| 32% | 76% \| 52% |
| **PW** | 52% \| 56% | 68% \| 48% | 96% \| 48% |

In order to evaluate the robustness of word-selective responses and their laterality across individual participants, we show individual Z-scores for the selected occipito-temporal EEG electrodes and gradiometers in Figure 6. For each participant, a larger blue than orange bar indicates a left-lateralised word-selective FPVS response. While the Z-scores decreased from CN to PW as before, and they did not exceed Z>2 for every participant, the majority of responses was left-lateralised, especially for gradiometers.

We further quantified the laterality of these responses. **Table 1** presents the statistical results for a comparison of Z-scores between the two hemispheres, for EEG, gradiometers above temporal scalp regions and for source estimates in the temporal lobes. For word-specific responses, gradiometers and source estimates produce reliably left-lateralised Z-scores for all three conditions. For EEG, laterality is marginally significant for the CN and NW conditions. For the base response, only source estimates in the CN condition show significant lateralisation, and in this case to the right. **Table 2** reports the percentage of participants that showed a numerically larger response in the left compared to the right hemisphere. The table also shows percentages for responses at the non-specific base frequency, as well as for source space results (see Figure 7). Except for EEG in the PW condition, all conditions produce left-lateralised word-selective responses in sensor space in the majority of participants. For gradiometers, the difference between word-selective and base responses is at least 25% in all conditions, suggesting that this lateralisation is not the result of unspecific features of our paradigm.

We present brain responses for ROIs in source space in the two separate hemispheres in Figure 7, with word-specific responses on the left and base frequency responses on the right. As apparent from the rightmost column in **Table 2**, at least 75% of participant show left-lateralised word-selective responses in all conditions. The PW condition is the most left-lateralised condition at 90%. In contrast, the base responses show left-lateralisation at or below 55%.

**Figure 6:**
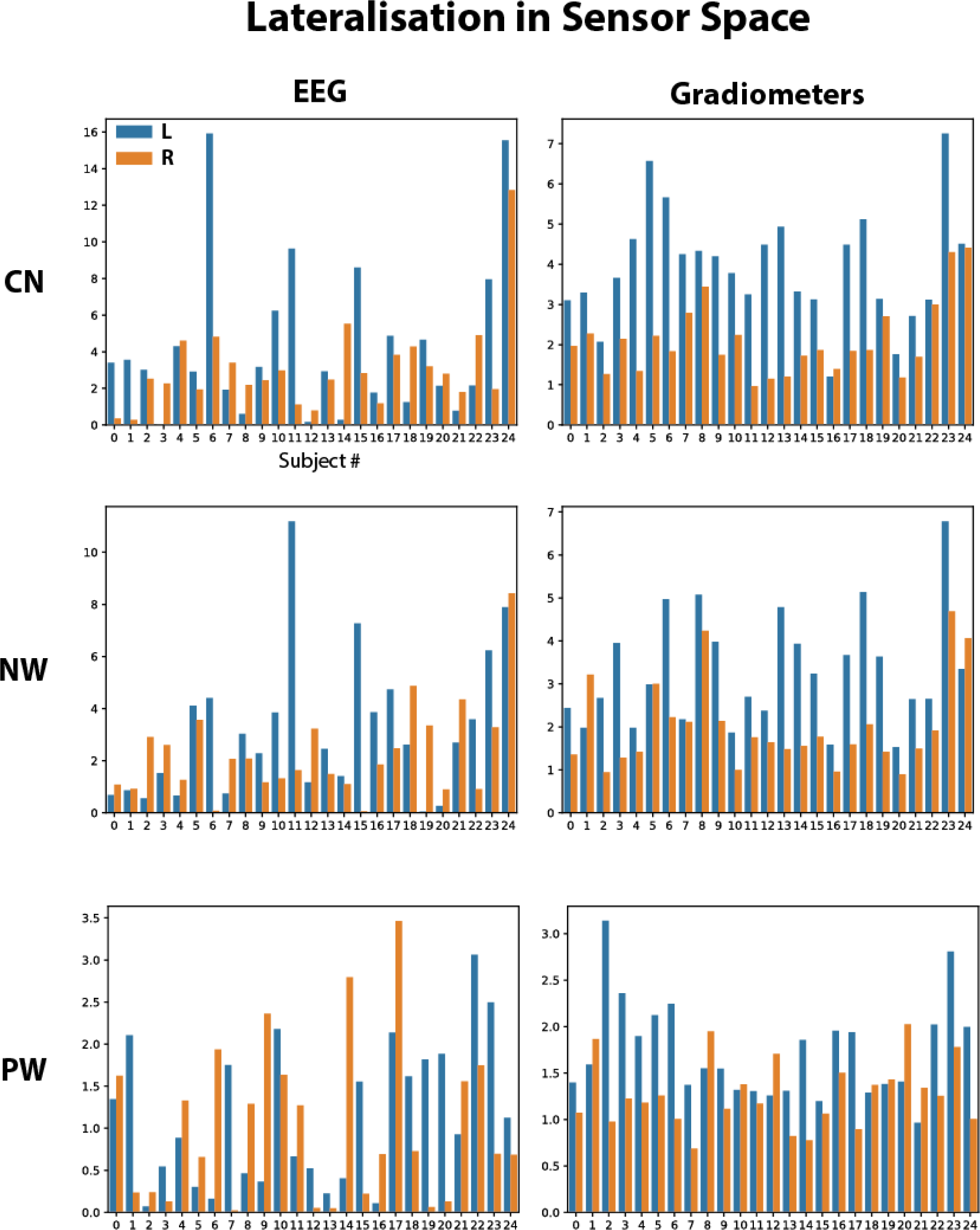
Z-scores for left and right sensor-space ROIs for individual participants. The bar graphs show average RMS Z-scores across left and right occipito-temporal EEG sensors (left) and temporal gradiometers (right), respectively. CN: consonant strings; NW: nonwords; PW: pseudowords.

**Figure 7:**
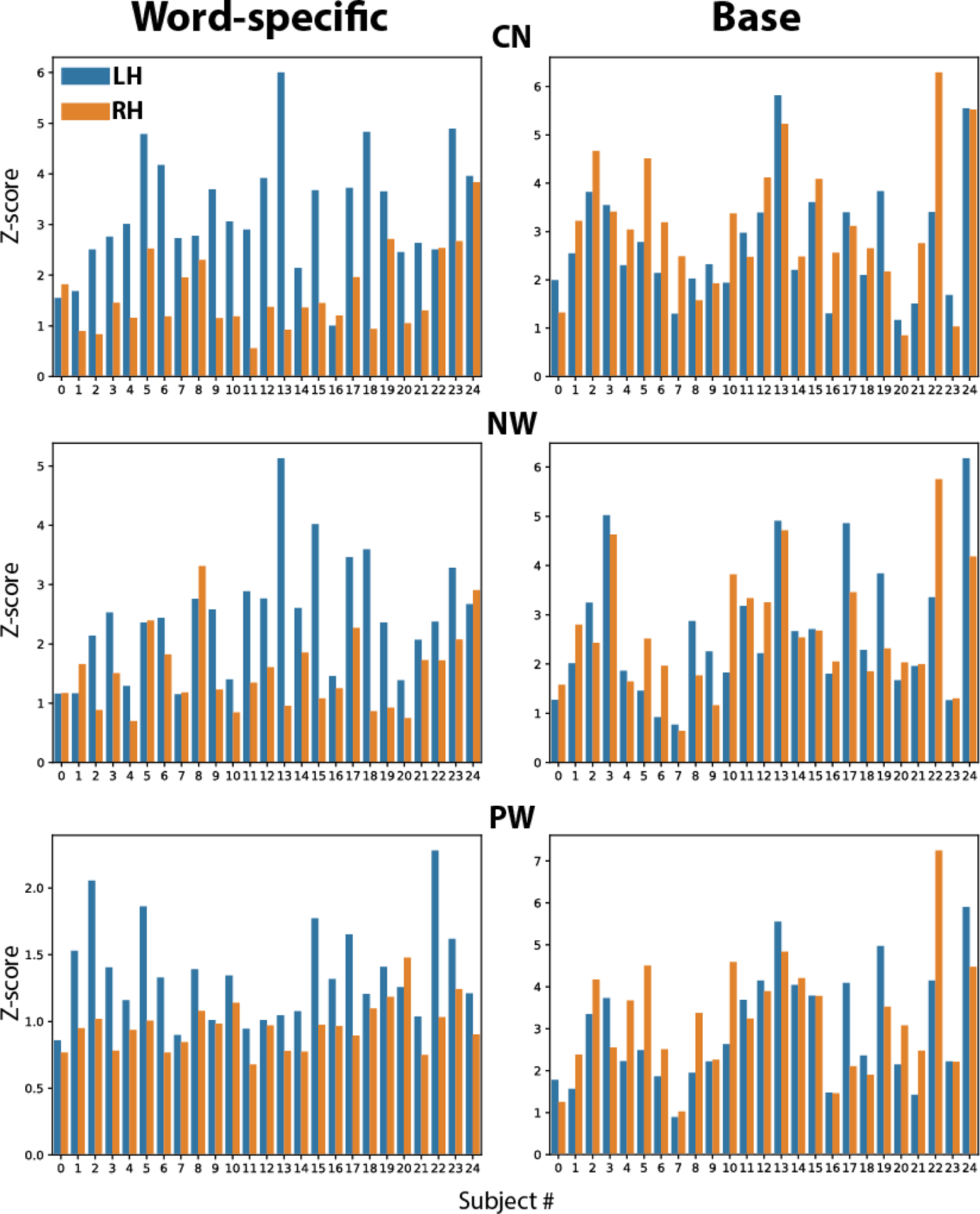
Z-scores for left and right temporal lobe ROIs for individual participants. The bar graphs show mean absolute Z-scores averaged across inferior, middle and superior temporal ROIs from the Desikan-Killiany cortical parcellation for individual participants (labelling as in Figures 6). Blue bars represent values for the left hemisphere (LH) and orange bars for the right hemisphere (RH), respectively. The left side shows bar graphs for word-specific responses and the right side for responses at the base frequency. CN: consonant strings; NW: nonwords; PW: pseudowords.

## Discussion

We used fast periodic visual stimulation (FPVS) with combined EEG and MEG to study rapid word discrimination processes in the human brain. Our results showed reliable and mostly left-lateralised word-selective responses in all conditions at the group level, even for the most fine-grained lexical contrast where words were embedded in orthographically-matched pseudowords. Word-selective responses were modulated by the word-likeness of base stimuli, as they were strongest when words were presented among consonant strings, weaker among nonwords, and weakest among pseudowords. Therefore, we extend for the first time the results originally obtained in French (Lochy et al., 2015) to the English language.

In the past decade, EEG coupled with FPVS has shown substantial advantages in terms of sensitivity, recording neural responses with a high SNR in just a few minutes, and objectivity, by identifying and quantifying the response at a pre-determined frequency, without any explicit task. The oddball FPVS design (reviewed in Rossion, Retter, & Liu-Shuang, 2020) measures neural responses to a deviant category of stimuli as an index of a *differential* processing between base and deviant stimuli (i.e., all common processes between base and deviant stimuli project to the common base rate response, which can be also objectively identified in the EEG spectrum). This method has been widely used among children learning to read (de Ghelcke et al., 2020, 2021; Lochy, Van Reybroeck, & Rossion, 2016; Lutz et al., 2024; Wang et al., 2024; Wang et al., 2023), in developmental disorders (e.g., dyslexia (Lochy, Collette, Rossion, & Schiltz, submitted)), and in healthy populations to study automatic lexical recognition (Lochy et al., 2024; Lochy et al., 2015; Marchive, Rossion, & Lochy, submitted) or semantic discrimination (Volfart, Rice, Lambon Ralph, & Rossion, 2021).

This approach provides important information about the neural bases of reading and left hemispheric lateralization. Previous FPVS-EEG studies have shown left occipito-temporal topographies for word recognition, modulated by the word discrimination contrast (Lochy et al., 2018; Lochy et al., 2024; Lochy et al., 2015) and reading abilities (Lochy et al., 2016; Wang et al., 2023) (Marchive et al. 2024). These findings are in agreement with the known brain regions implicated in reading and visual word recognition, such as the left ventral occipito-temporal cortex (lateral fusiform gyrus/occipito-temporal sulcus) (Cattinelli, Borghese, Gallucci, & Paulesu, 2013; Dehaene, Cohen, Sigman, & Vinckier, 2005; Schurz et al., 2015; Wandell & Le, 2017), and align with fMRI studies showing left dominance in the VOTC for pre-lexical or lexical processes (Jobard, Crivello, & Tzourio-Mazoyer, 2003; Vinckier et al., 2007), notably in the visual word form area (VWFA; (Cohen et al., 2000; Cohen et al., 2002; Dehaene et al., 2005; Dehaene et al., 2010). In the context of our FPVS results, it is important to note that fMRI studies of word recognition invariably show some degree of bilateral yet asymmetric vOT responses to written words and letter strings (Behrmann & Plaut, 2015), a pattern that is mirrored in case-series studies of patients with damage to these regions (Behrmann & Plaut, 2014; Roberts et al., 2013).

Thus, an approach involving implicit, rapid and objective measurement is important to extend to other languages. Although word-selective responses were found in German (Aristei, Lochy, Rossion, & Schiltz, 2017), a recent study encountered difficulties replicating these effects in ten English participants (Barnes et al., 2021). Lochy et al. (2024) discussed this discrepancy as potentially stemming from linguistic differences between French and English in orthographic neighbourhood density (N), or methodological differences in the choice of pseudowords as highlighted in the Introduction.

In the current study with a larger number of participants, we designed pseudowords matched pairwise to the word stimuli for consonant/vowel structure, bigram frequency, and neighbourhood density. We successfully extended results from the French studies (Lochy et al., 2024; Lochy et al., 2015) to English. First, we found a word-selective response for all conditions at the group level, modulated by the wordlikeness of base stimuli (mirroring behavioural effects found previously in behavioural lexical decision studies(Evans et al., 2012)). As explained by Lochy et al. (2015), the word discrimination amplitude depends on the nature of the list context. For example, when words are discriminated among pseudo-letters, the response is larger and more bilateral (although with a left hemispheric dominance) than for words among nonwords (NW) or pseudowords (PW) (Lochy et al., 2015). Indeed, discriminated against pseudo-letters, responses to written words may reflect not only lexical and orthographic processes but also, more basically, the contrast of real letters to non-existing letter-like forms, thus generating more bilateral, lower-level, visual responses. Similarly, words among NW appear to trigger a larger amplitude response than words among PW (Lochy et al., 2015), possibly because this coarser contrast may be based on pre-lexical orthographic plausibility detection by comparison to the finer contrast. Here, our EEG results also evidence this modulation of response strength, with stronger responses for words among consonant strings, than among non-words or pseudowords. Importantly, word-selective responses at the group level were located over the left vOTC. Moreover, besides extending the results from (Lochy et al., 2015) to the English language, we also presented MEG in addition to previous EEG results, and used their combination for analyses in source space. Only one previous study used EEG and MEG with an FPVS paradigm, on face categorization (Hauk et al., 2021), and provided useful information on distributed source estimation over left and right hemispheres. Here, our source space results complemented those in sensor space. Word-selective responses in the group analysis were plausibly localised to the temporal lobe and left-lateralised. As in sensor space, they were most reliable for words in consonant string context, followed by nonwords and pseudowords. Significant Z scores spread along the left temporal lobe for words embedded in consonant string showing a coarser discrimination contrast with a more bilateral response over the occipital visual areas. More focal and left-lateralized responses appear with finer-grained lexical discrimination. However, the differences in SNR ratio between conditions makes a more detailed comparison of source distributions difficult. Nevertheless, the peaks of these distributions in the left temporal lobe were in similar locations across conditions, suggesting that the underlying distributions are similar and mainly differ with respect to their noise levels (see also Lochy et al., 2018).

Second, we were able to assess individual discrimination responses. Previous studies on face-selective FPVS responses suggested that reliable category-specific responses could be obtained for the majority of participants with only a few minutes of data recording, raising the possibility that FPVS could become a standard tool for localiser scans or to monitor perceptual and cognition processes in clinical populations (Lochy et al., 2015; Rossion et al., 2015). With respect to word-like stimuli, the study by Lochy et al. (2015) reported that word-specific responses for French words among well-matched pseudowords could be detected as above baseline for the majority of participants, especially when topographies and laterality were taken into account. This high rate of response detection at the individual level was recently replicated (Lochy et al., 2024). Such a high sensitivity is an extremely important factor for future investigation of clinical populations (e.g. dementia) and developmental disorders (e.g., dyslexia) or for assessing the emergence of brain specialization for print and lexical representations in young children. Interestingly, in the present study MEG and source estimation produced a more consistent pattern with respect to lateralisation of word-selective responses than EEG. While EEG produced left-lateralised responses for more than 50% of participants in the consonant and non-word conditions, it was exactly 50% in pseudoword context. In contrast, gradiometers achieved more than 65% and 75%, respectively, and for pseudowords lateralisation in source space reached its highest level at 90%. Future studies will reveal whether MEG, possibly in combination with EEG, can reveal word-specific brain responses at an individual-participant level. New developments in non-cryogenic on-scalp MEG sensors, such as optically pumped magnetometers (OPMs), which might achieve higher signal-to-noise ratios than conventional MEG systems, are particularly promising in this respect (Boto et al., 2018).

## Acknowledgments

The project was partly funded by a *Lorraine Université d’Excellence* (LUE) grant to foster international collaborations between Université de Lorraine and University of Luxembourg (UL_IRP 2022). AL is supported by the Fonds National de la Recherche du Luxembourg (FNR-CORE C21/SC/16241557/READINGBRAIN). MALR and OH are supported by intra-mural funding from the Medical Research Council (UK: MC_UU_00030/9). MM is supported by Lorraine Université d’Excellence with a DrEAM grant. For the purpose of open access, the author has applied a CC BY public copy-right license to any Author Accepted Manuscript version arising from this submission.

## Author contributions

All authors were involved in conceptualising this study. AL, MM, BR, MALR created the stimuli and designed the experiment. MM and OH acquired the data. OH analysed the data. All authors were involved in discussing the results. OH wrote the first draft of the manuscript. All authors contributed to the reviewing and editing of the manuscript.

## Ethics Statement

This study was approved by the Cambridge Psychology Research Ethics Committee.

## Data and Code Availability

The data are available from the MRC Cognition And Brain Sciences’ data repository on request. The code used to analyse the EEG/MEG data is openly available on github: https://github.com/olafhauk/FPVS_WORDS.

